# Genomic data confirms phenotypic predictions of hybridization between cryptic Hawaiian cricket species

**DOI:** 10.1101/2025.09.09.675270

**Authors:** Raunak Sen, Kerry L. Shaw

## Abstract

Hybridization is increasingly recognized as an important driver of species radiations, yet its detection in morphologically cryptic taxa remains challenging. The Hawaiian crickets of the genus *Laupala* represent a rapid non-adaptive radiation in which species are morphologically indistinguishable but differ in acoustic mating signals. Here we report the first discovery of natural hybridization in the *Laupala* radiation. During field surveys on Maui, we observed crickets with intermediate pulse rates and unusually high acoustic variation at sites where the ranges of *Laupala makaio* (slow singer) and *Laupala orientalis* (fast singer) converge. We hypothesized these represented natural hybrids and tested this prediction using genomic data from genotyping-by-sequencing (GBS). Maximum likelihood phylogenetic analysis showed that fast-singing putative hybrids clustered with *L. orientalis* and the slow-singing ones with *L. makaio*, yet all putative hybrids formed a cohesive geographic group. Using 9,345 ancestry-informative markers, we calculated hybrid indices and heterozygosity values for triangle plot analysis. All hybrids showed excess *L. makaio* ancestry (predicted from analysis of pulse rate patterns) and were classified as later-generation hybrids or backcrosses to *L. makaio*, with no F1 hybrids detected. Critically, we found a strong positive correlation (R = +0.74, p = 4.1×10⁻⁶) between pulse rate and hybrid index at the site with highest pulse rate variation, showing that genomic ancestry predicts pulse rate phenotype. Morphological comparisons confirmed that hybrids were cryptic, reinforcing the difficulty of detecting hybridization without genomic data. Together, our results provide the first demonstration of natural hybridization in *Laupala*, showing that phenotypic predictions based on acoustic intermediacy can be confirmed with genomic evidence. The discovery of natural hybridization in *Laupala* opens new avenues for investigating how hybridization contributes to rapid diversification in non-adaptive radiations, paralleling patterns observed in some adaptive radiations.

## Introduction

Species radiations - the rapid diversification of a lineage into multiple species exhibiting phenotypic and genotypic diversity, represent some of the most dramatic examples of evolutionary diversification in nature. These radiations can be “adaptive” if the species exploit different ecological niches and ecology seems to play an important role in their diversification (Schluter, 2000), as in the case of Darwin’s finches (P. R. Grant & Grant, 2020), African crater lakes cichlids (Seehausen, 2006), *Heliconius* butterflies (Jiggins, 2008), Caribbean *Anoles* lizards (J. Losos, 2009) and Hawaiian *Drosophila* (Craddock & Kambysellis, 1997). Or they can be “non-adaptive” if the species are ecologically cryptic and the role of ecology in the speciation process is unclear (Gittenberger, 1991). Some examples of non-adaptive radiations include many clades of molluscs (Gittenberger, 1991; Sauer & Hausdorf, 2009; Wilke et al., 2010), some reptile species (K. H. Kozak et al., 2005; Lobón-Rovira et al., 2024; J. B. Losos & Miles, 2002), grasshoppers (Mayer et al., 2010) and Hawaiian crickets (Shaw, 1995). These explosive bursts of speciation have captivated evolutionary biologists for decades and provide natural laboratories for understanding the mechanisms driving diversity. Studying species radiations help us understand the extrinsic ecological factors, intrinsic lineage specific biological processes, or an interaction between the two, which aids rapid speciation. Despite extensive research, a fundamental question remains: why do some lineages radiate while others do not? (De-Kayne et al., 2025)

Often species radiation have both an adaptive and non adaptive component as various processes might be fueling rapid speciation and each process might be more relevant at different timepoints during the radiation (Schön & Martens, 2004). Many empirical studies have highlighted some potential drivers of radiation: hybridization and genetic admixture between divergent lineages (De-Kayne et al., 2022; Meier et al., 2017), sexual selection (Mendelson & Shaw, 2005; Sauer & Hausdorf, 2009; Wagner et al., 2012) and the acquisition of key innovations (Hunter, 1998; Schluter, 2000). Hybridization, in particular, is emerging as an important process fueling species radiations because it is not constrained by genetic variation, in contrast to other lineage specific factors. In fact hybridization creates new genetic variants and novel phenotypes, thereby increasing adaptive potential (B. R. Grant & Grant, 2008). The shuffling of ancient polymorphisms via hybridization has fueled the adaptive radiations of African cichlids (Meier et al., 2017), *Heliconius* butterflies (K. M. Kozak et al., 2021) and Hawaiian *Metrosideros* plants (Choi et al., 2021). Novel phenotypes created through hybridization can colonize new ecological niches, aiding further diversification; for e.g., *Helianthus* sunflower hybrids have combined additive effects of parental loci for salt tolerance, colonizing habitats which neither parental species could (Lexer et al., 2003). Hybrid speciation events can also rapidly increase species diversity (Lamichhaney et al., 2018; Rosser et al., 2024). Various hypotheses have been proposed to better understand the mechanisms in which hybridization can promote explosive speciation rates (Marques et al., 2019; Seehausen, 2004, 2013).

Demonstrating the role of hybridization in promoting species radiations requires first establishing that natural hybridization actually occurs within these rapidly diversifying lineages. This has been shown in many lineages, particularly helped by the observation of visually conspicuous phenotypic intermediates or recombinants, as in body size and beak shape in Darwin’s finches (P. R. Grant & Grant, 2000) or recombinant wing patterns in Heliconius butterflies (Gilbert, 2003). However, detection of hybridization becomes considerably more challenging in cryptic species complexes - groups where species are morphologically indistinguishable to the human eye but are genetically distinct (Bickford et al., 2007; Fišer et al., 2018). Studies suggest that hybridization in cryptic species is significantly underrepresented in the literature, creating an ascertainment bias in our understanding of hybridization patterns (Maguilla & Escudero, 2016; Mallet, 2005; Quattrini et al., 2019). In cryptic species, hybrid individuals are often difficult to detect because they have intermediate phenotypes which are subtle or expressed in non-visual traits such as acoustic signals or chemical cues that require specialized techniques to measure (Keller & Carl Gerhardt, 2001; J. Purcell et al., 2016). Consequently, many potential cases of hybridization may remain undetected, particularly in diverse but understudied groups such as invertebrates.

The advent of next-generation sequencing and other genomic tools and analyses has revolutionized our ability to detect hybridization in cryptic species (Taylor & Larson, 2019; Twyford & Ennos, 2012). High-throughput sequencing approaches which include reduced representation sequencing (RAD-seq, ddRAD, GBS), whole-genome sequencing and transcriptomics have enabled researchers to identify genomic signatures of hybridization even in the absence of clear morphological intermediates (Belintani et al., 2023; Herrera-Aguilar et al., 2009; McKenzie et al., 2021; Patel et al., 2015; Quattrini et al., 2019; Slager et al., 2020; vonHoldt et al., 2016). These genomic tools allow for the detection of hybridization through estimation of phylogenies, analyses of allele frequencies, ancestry blocks and introgression patterns across the genome (Gompert et al., 2017; Twyford & Ennos, 2012). Genomic approaches have revealed that hybridization is more common across the tree of life than previously thought (Dagilis et al., 2021; Goulet et al., 2017; Mallet et al., 2016; Ottenburghs, 2023; Shurtliff, 2013; Taylor & Larson, 2019).

The non-adaptive radiation of the Hawaiian crickets of the genus *Laupala* (Orthoptera: Gryllidae: Trigonidiinae: Laupala) represent an ideal system for investigating the potential role of hybridization in fueling species radiations. This explosive radiation contains 38 species distributed across different islands of the Hawaiian archipelago and has the highest known rate of speciation in invertebrates (Mendelson & Shaw, 2005). Species are morphologically and ecologically cryptic but differ in male songs and female preferences for those songs (Otte, 1994; Shaw, 1999; Shaw & Herlihy, 2000). Males of different *Laupala* species sing species-specific songs that differ primarily in their pulse rate - the number of pulses produced per second.

Females prefer conspecific songs over heterospecific songs in lab phonotaxis trials. Some species can be successfully crossed in the lab. Lab hybrid males have intermediate pulse rates of their parental species and hybrid females prefer these intermediate pulse rates (Blankers et al., 2019; Shaw, 2000b). Male song and female preference have co-evolved (Grace & Shaw, 2011), primarily due to genetic coupling (Xu & Shaw, 2021). Together, male song and female preference is an important pre-mating barrier to gene exchange, and this genus is model clade to investigate the link between divergence of mating traits and speciation.

Here we report the first discovery of natural hybridization in the *Laupala* radiation. Although different species can be hybridized in the lab, we have not discovered any natural hybrids in the wild until now. This could be because the species are morphologically indistinguishable from one another and a non-visual trait (acoustic) expressed by only one sex (males) and only during certain time points, distinguishes them. We noticed a huge variation in pulse rate in a population of crickets in the island of Maui in Hawaii. Other nearby populations had a pulse rate not heard in any *Laupala* species on the island. On formal analysis of the pulse rate patterns and armed with the knowledge of the pulse rates of the *Laupala* species on the island, we hypothesized that the individuals were hybrids between morphologically indistinguishable *Laupala makaio* and *Laupala orientalis* (see **Figure 1**), which had parapatric distributions at the site of hybridization. Between these two species *Laupala makaio* is the “slow singer” which sings at 0.6 pulses per second (pps) and *Laupala orientalis* sings at 2.2 pps (see **Figure 2**), making it the “fast singer”. The putative hybrid individuals, which are also morphologically similar to the parental species, had intermediate pulse rates which can be explained by hybridization. Based on previous knowledge on the inheritance patterns of pulse rate from creating lab hybrids (Blankers et al., 2019; Shaw, 2000b), we knew that *Laupala* hybrids have an intermediate pulse rate of their parental species. Thus, we hypothesized that the observation of intermediate pulse rates in the wild is due to natural hybridization between *Laupala makaio* and *Laupala orientalis*. The current (for *L. makaio*) and historical (for *L. orientalis*) geographical proximity to the site where the putative hybrids were observed also bolstered our hypothesis. We tested this hypothesis of hybridization using genomic analyses.

**Figure 1:**
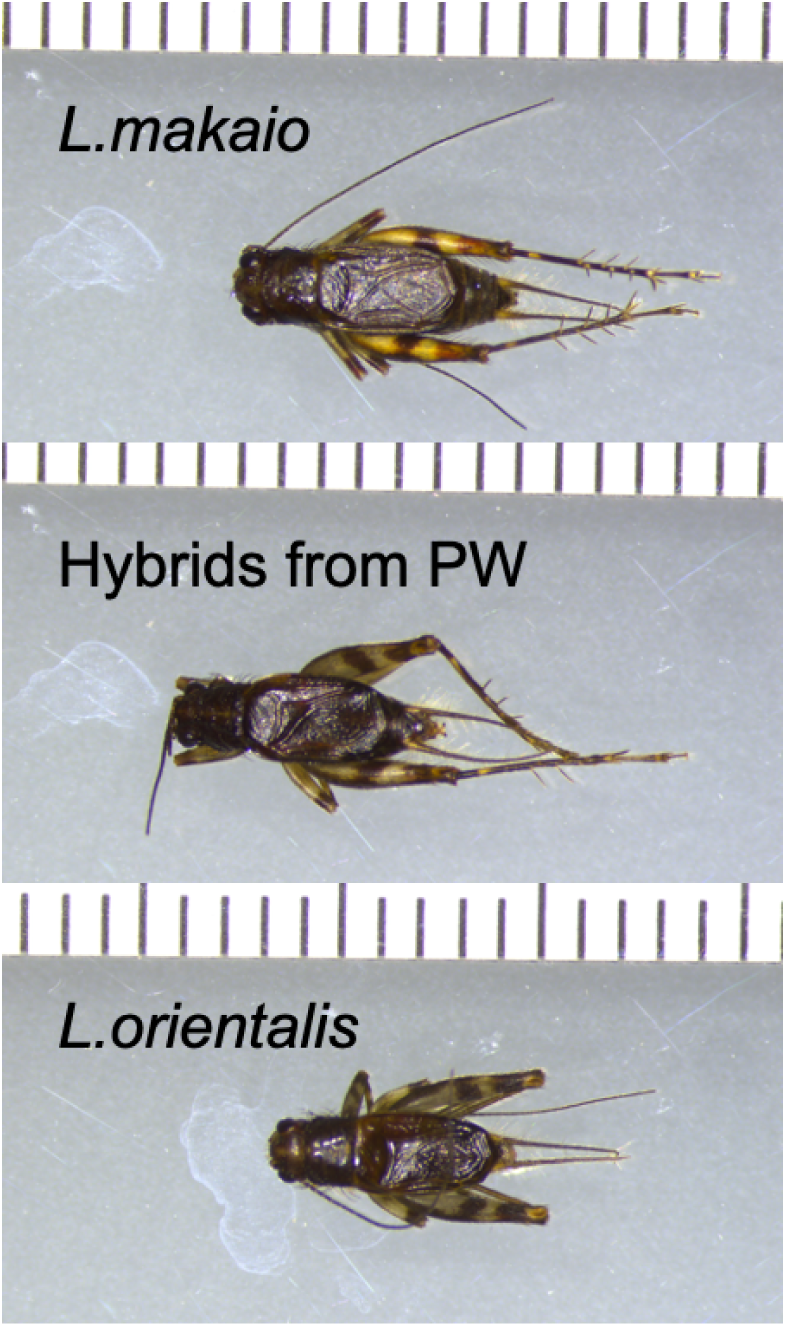
(from top to bottom) Dorsal view of a male cricket of *L. makaio*, hybrids from PW and *L. orientalis* (respectively) under the same magnification and lighting conditions, shows how morphologically indistinguishable they are.

**Figure 2:**
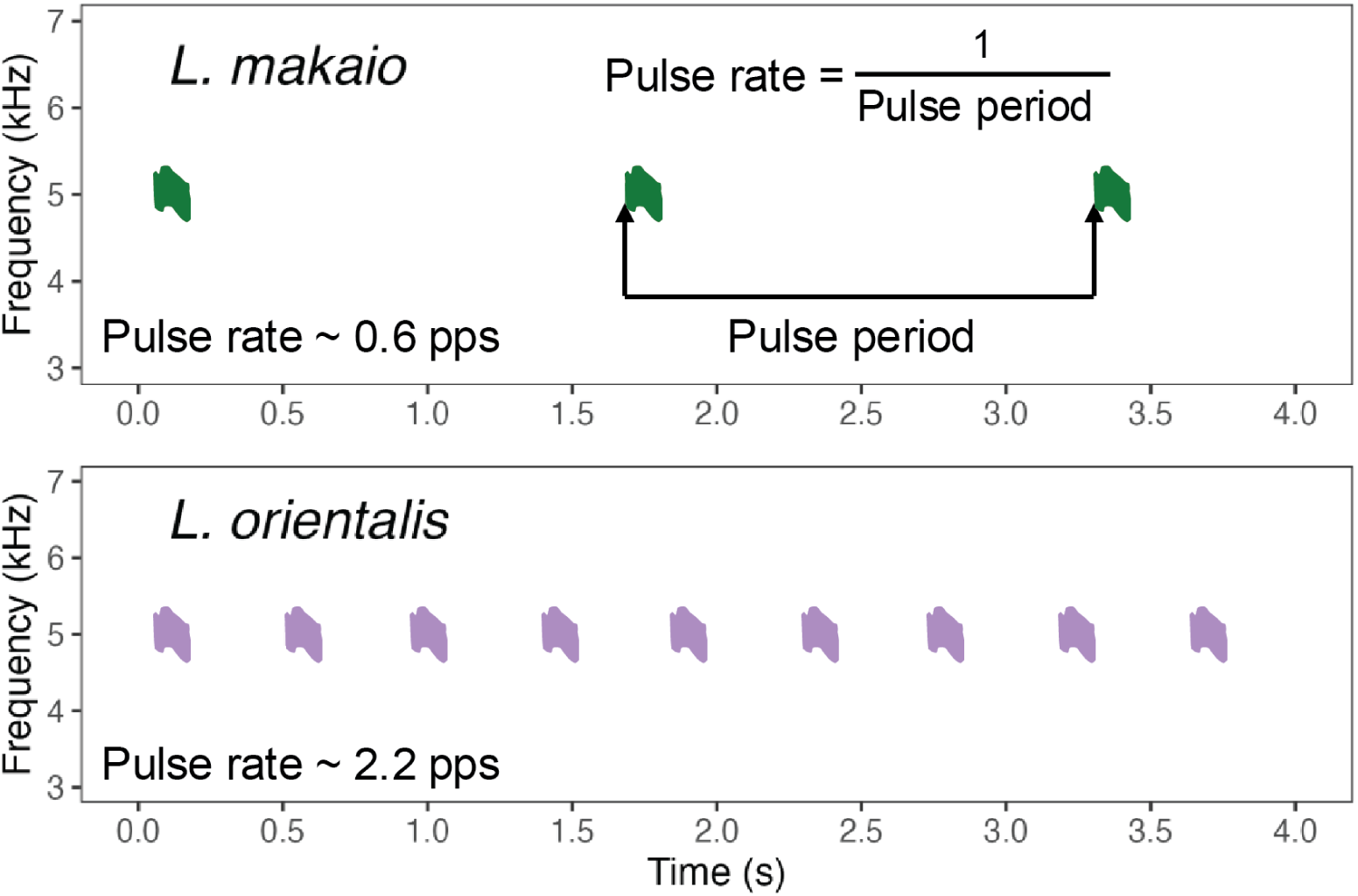
A schematic of a sonogram of *Laupala* cricket song, which is a simple train of pulses. The distance between the beginning of two consecutive pulses is the pulse period. The inverse of it is the pulse rate. In our case, between the two hybridizing species, *L. makaio* (top) is the “slow singer”, singing at 0.6 pulses per second (pps) and *L. orientalis* (bottom) is the “fast singer”, singing at 2.2 pps.

We extracted DNA from the wild collected individuals and generated a single nucleotide polymorphism (SNP) dataset by employing GBS (a reduced-representation sequencing approach)(Elshire et al., 2011). This SNP dataset was then analyzed to look at population structure and estimate phylogenies. A common goal of hybridization studies is to classify individuals into different classes of hybrids (F1, F2, backcross or later-generation hybrids) or as parental species (Fitzpatrick, 2012; Gompert & Buerkle, 2013). To that end, we extracted ancestry informative markers (AIMs) to distinguish between the two parental species and calculated the hybrid index and heterozygosity of each hybrid. The relationship between hybrid index and heterozygosity was visualized by making a Triangle plot. Triangle plots are powerful methods to infer admixture (Wiens & Colella, 2025). Using the Triangle plot, we classified the hybrids into different classes. Such a classification is often a starting point to generate hypotheses of reproductive isolation, selection and introgression in a hybrid zone. Based on the observation that the pulse rate of the hybrids were closer to that of *L. makaio*, the analysis of pulse rate patterns of hybrids and the knowledge of inheritance patterns of pulse rates, we predicted that (i) all hybrids have a higher ancestry of *L. makaio* compared to *L. orientalis* (ii) most hybrids were later generation hybrids or back-crosses into *L. makaio*. We also tested for phenotype-genotype correlations between pulse rate and hybrid index (or ancestry) of hybrids at PW (the site with the highest variation in pulse rate). If the variation of pulse rate at PW is a result of hybridization, we would predict a positive and significant correlation of pulse rate and hybrid index (in our case, ancestry of the faster species *L. orientalis*). In other words, we predict that hybrids with a higher ancestry of *L. orientalis* in their genome will sing at faster pulse rates.

## Methods

### Collection of wild cricket samples

We collected wild crickets from five sites in May 2021 and May 2023 on the island of Maui, state of Hawaii, USA. These sites were located on different elevations on a mountain called “Palikea Peak”, on the South-Eastern part of the island (**Figure 3**). The sites were Summit Area (SA), Pig Wallow (PW), Lunch Spot (LS), Old Koa (OK) and Paper Bark (PB). SA is the parental range of *Laupala makaio* and other four sites had crickets with varying levels of song intermediacy. For genomic data collection, some historical samples were also included. These were collected from the “Dog leg” (DL) site at Palikea Peak (also the range of *Laupala makaio*) and a nearby site called “Kipahulu” (KP) (historical range of *Laupala orientalis*, currently non-existent). For outgroup comparisons, we collected a distantly related co-occurring species *Laupala eukolea* from Palikea Peak and also used some historical samples from Kipahulu. Some samples from another distant (from Palikea Peak) population of *Laupala orientalis* (from Makapipi; MP) were also included in the sequencing dataset but they were not used for any downstream analysis reported in this paper because we found they were not the population that was hybridizing. A map of all sampling locations is provided in **Figure 3**. A table containing the GPS coordinates, elevation of each site and the number of crickets sequenced from that site is given in Supplementary Information Table S1. Juvenile crickets were transported alive from Hawaii to our lab in Ithaca, NY and were raised until maturity.

**Figure 3:**
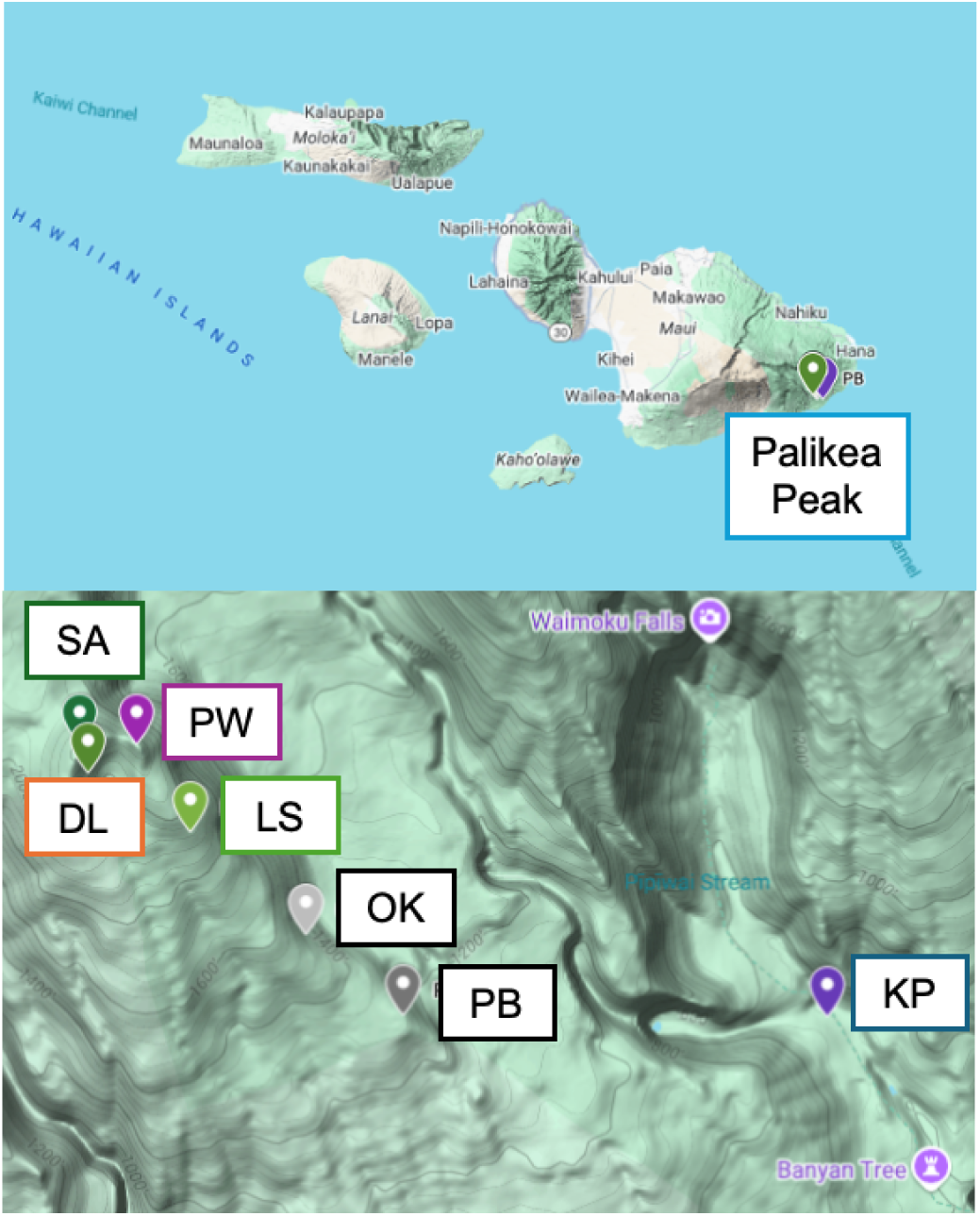
(top) Map of the sampling location (Palikea Peak) on the East side of the island of Maui in Hawaii (bottom) Map of the different sampling sites where *L. makaio* (DL and SA) and the hybrids (PW, LS, OK and PB) are found and the site (KP) where historically *L. orientalis* occurred and was collected.

### Song data collection and analysis

For all the crickets collected in the wild in 2021 and 2023, we recorded the songs of mature male crickets with an Olympus WS-852 digital recorder in a temperature-controlled room (19-21 C). The sound files were analyzed using RavenLite 2.0. Pulse period was measured as the time difference between the start of two consecutive pulses (as shown in **Figure 3**). The mean pulse period was calculated from five independent measurements from a single pulse train. We calculated the pulse rate (in pulses per second or pps) by taking the inverse of the mean pulse period. In our pulse rate histogram (see **Figure 4**), we also included previous data (Shaw, 2000a) on pulse rates recorded in the field for *L. orientalis* (from site KP), for comparison.

**Figure 4:**
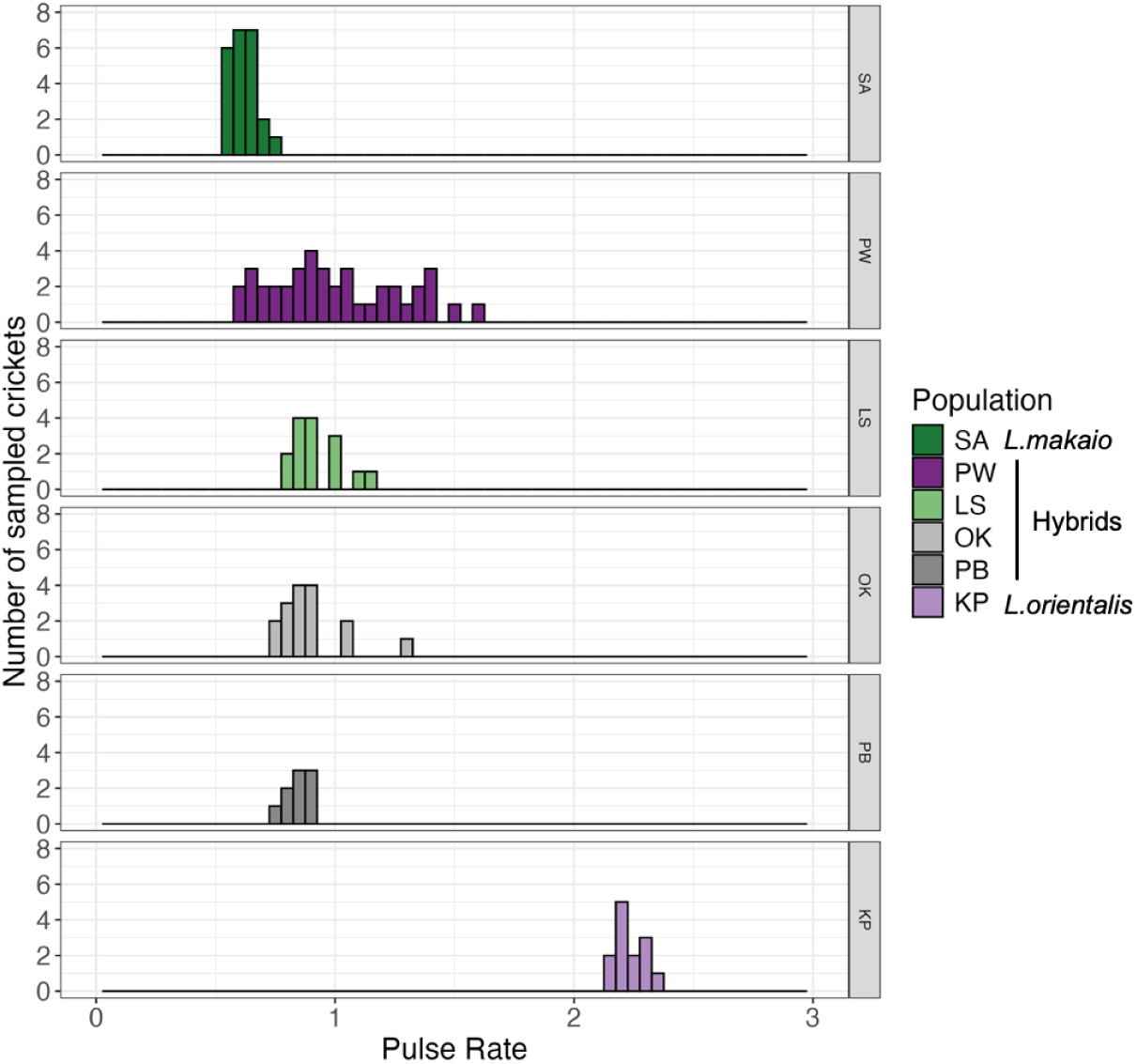
Histogram of the distribution of pulse rates of crickets collected from each population. PW has the highest variation in pulse rate compared to other populations.

### Morphology data collection and analysis

From crickets, we dissected and imaged hindlegs, wings and ovipositors (only for females) under a microscope with an appropriate scale. We processed the images in ImageJ and measured femur length, forewing length and ovipositor length (only in females). Only a subset of the cricket samples were measured for morphology: hybrids from the site of maximum pulse rate variation (PW) (See Results), slow singing parental species *L. makaio* from SA and fast singing parental species *L. orientalis* from MP (n=20 per group). We used two-tailed Student t tests, ANOVA and Tukey’s post hoc HSD to test for significance between groups. All statistical tests were performed in RStudio Version 2024.09.1+394.

### DNA extraction and sequencing

DNA was extracted from the crickets post dissection with the QIAGEN DNeasy Blood and Tissue Kit with three minor changes to the standard manufacturer’s protocol of extracting DNA from insects: (i) we added double the amount of Proteinase K in the first incubation step (ii) increased the time for first incubation to 12-16 hrs and (iii) we repeated the last elution step with 50 μL AL. We measured the concentration of the extracted genomic DNA using a Qubit and checked its quality by running agarose gels. Samples with predominantly high molecular weight DNA and with a concentration above 10 ng/μL were sent to the University of Wisconsin Biotechnology Center for library preparation and sequencing for Genotype by Sequencing (GBS)(Elshire et al., 2011). Briefly, the enzyme cutters `MspI` and `PstI` were used to digest DNA, which was then ligated to barcodes and common Illumina TruSeq adapters. The ligated DNA was then amplified using PCR and pooled for sequencing. Libraries were sequenced on half a lane of a Illumina NovaSeq X Plus, which produced approximately 450M paired end reads of insert size length of 150 bp.

### Genomic data filtering

Sequenced reads were demultiplexed using `process_radtags` from Stacks 2.67 (Rochette et al., 2019) with the `--force-poly-g-check` flag, because there is an overrepresentation of poly-G sequences in data generated by a NovaSeq machine (De-Kayne et al., 2021). We checked the demultiplexed reads of a few samples with `FastQC` (Simon, 2010) and found that they contained Illumina adapters and some poly-G sequence remnants which were removed using `Trimmomatic v 0.39` (Bolger et al., 2014) in ‘palindrome mode’. We used these settings for trimming: `ILLUMINACLIP:TruSeq2-PE.fa:2:30:10:2`. We also used `Trimmomatic` to trim reads for quality using the following setting: `LEADING:3 TRAILING:3 SLIDINGWINDOW:4:15 MINLEN:36.`Trimmed reads were then aligned to the *Laupala kohalensis* genome assembly (Blankers et al., 2018) using `bwa-mem2 version 2.2.1` (Li & Durbin, 2009; Vasimuddin et al., 2019) to make `sam` files. We used `samtools v 1.2` (Li et al., 2009) to convert the files from `sam` to `bam`, sort it and then filter by mapping quality (MQ) score to remove reads with MQ < 20. The filtered `bam` files were used to call variants using `bcftools v 1.20` (Li, 2011) to create a VCF file. We then filtered the VCF file using `vcftools` (Danecek et al., 2011) to remove any loci with more than 20% of the data missing and any in-dels, retaining only single nucleotide polymorphisms (SNPs). A third round of filtering was done (again using `vcftools`) to only retain the SNPs which have a `phred` quality score > 40 (high quality base calls), bi-allelic, have a minimum average read depth of 10 and maximum 40 and a minor allele count > 1. Seven samples were removed because they did not pass the filtering steps. We retained 185 samples and 546,852 SNPs.

### Population structure analysis

We conducted a principal component analysis (PCA) with genome-wide SNPs to assess the population genetic structure of the crickets at Palikea Peak (which includes the site DL and SA where *L. makaio* occurs and other sites where the hybrid individuals occur) and *L. orientalis* at KP. A genomic PCA summarizes the major axes of variation in allele frequencies which are independent of each other. The VCF file was subsampled to only include the above-mentioned samples. We performed our PCA in `plink v 1.90b7` (Chang et al., 2015; S. Purcell et al., 2007). To fulfil one of the major assumptions of a PCA i.e. there should be no spurious correlations between the measured variables, we pruned our dataset of variants which are in linkage. That left us with 157,419 SNPs which were used to make the PCA. We plotted the PCA using custom R scripts (adapted and modified from https://speciationgenomics.github.io/). We did not make a standard STRUCTURE or ADMIXTURE plot because these model-based population structure analyses do not take spatial auto-correlation into account and our data had a significant signal of genetic Isolation by Distance (IBD) (see Supplementary information Table S2 and Figure S1).

### Estimation of Phylogeny

We made a phylogeny to test our hypothesis of hybridization. We selected 5 samples of *L. makaio* (randomly 2 from DL and 3 from SA), the fastest and slowest five singers from the hybrid population at PW (total 10 from PW), and 5 samples of *L. orientalis* (randomly from KP). Five *L.eukolea* samples were included as an outgroup. A total of 25 individuals were used to make the phylogeny. If the crickets at PW are an outcome of hybridization between *L. makaio* and *L. orientalis*, we would expect the slower singers to be more closely related to *L. makaio* and the faster singers to be more closely related to *L. orientalis* in a phylogeny. However, as all the individuals at PW (both slow and fast singers) are at the same geographical location, they are probably exchanging genes, so they would be closely related to each other. The filtered VCF file was subsampled to only include the above-mentioned samples. After linkage pruning using `plink v 1.90b7` (Chang et al., 2015; S. Purcell et al., 2007), the VCF file was converted to the PHYLIP format suitable for phylogenetic programs using `vcf2phylip` (Ortiz, 2019). We used the PHYLIP file to make a phylogeny in `PhyML 3.0` (Guindon et al., 2010) by submitting the file in their “Online execution” platform (http://www.atgc-montpellier.fr/phyml/) and used standard settings. 48,164 SNPs were used to build the phylogeny.

### Analysis of hybridization by Triangle Plots

A hybrid index vs heterozygosity plot, often known as a triangle plot, was built using the package `triangulaR` (Wiens et al., 2025). From the filtered SNP dataset used in population structure analysis, we extracted ancestry informative markers (AIMs) which can distinguish between the two parental species. The SA population of *L. makaio* and the KP population of *L. orientalis* were chosen as the two parental populations. The DL population was not chosen for *L. makaio* because of low sample size (n=4) which is not ideal for calculating AIMs. Alleles with a frequency difference > 0.75 between the two parental species were used as AIMs because simulation studies have shown that alleles with frequency differences above this value strike an optimal balance between being ancestry informative and providing sufficient markers for precise hybrid index and heterozygosity calculations, particularly in later-generation hybrids (Wiens et al., 2025). This threshold yielded 9345 AIMs which were used to calculate the hybrid index and heterozygosity of each individual. The relationship between hybrid index and heterozygosity was visualized in a Triangle plot. Following Scordato et al., 2017, we classified individuals into different hybrid classes: crickets with hybrid indices ranging from 0.02 to 0.25 or 0.75 to 0.98 were considered backcrosses to one or the other parental type. Crickets with hybrid indices >0.25 and <0.75 and heterozygosity ≥ 0.80 were classified as F1 hybrids. Crickets with these intermediate hybrid indices but lower heterozygosity (<0.80) were considered later-generation hybrids. Although it is very difficult to classify individuals with complex ancestry, this approach provides a rough summary of the distribution of genomic admixture.

### Phenotype Genotype Correlations

We performed a Pearson Correlation test to test for association between pulse rate and hybrid index for n=29 males at PW. We calculated the correlation coefficient (R) and the p-value to assess the statistical significance of the association.

## Results

### Analysis of pulse rate data

The distribution of pulse rate at each population is shown as a histogram in **Figure 4**. The mean and standard deviation (SD) of the pulse rate of male crickets at each site is given in **Table 1**.

**Table 1:** Sample size, mean and standard deviation (SD) of pulse rate (in pulses per second or pps) of male crickets at each site.We should note that these are sample sizes of male crickets only, as only males can sing.

| Population | Species | Sample size | Mean (in pps) | SD (in pps) | Sampling year |
| --- | --- | --- | --- | --- | --- |
| SA | *L. makaio* | 23 | 0.62 | 0.05 | 2021 and 2023 |
| PW | Hybrids | 29 | 1.01 | **0.27** | 2021 and 2023 |
| LS | Hybrids | 15 | 0.92 | 0.1 | 2021 and 2023 |
| OK | Hybrids | 16 | 0.9 | 0.14 | 2021 |
| PB | Hybrids | 9 | 0.85 | 0.05 | 2021 and 2023 |
| KP | *L. orientalis* | 13 | 2.24 | 0.07 | 2000 and 2001 |

As we can see from **Table 1**, the crickets at PW have the highest SD (and thus variance) of pulse rate, which is statistically significant compared to other populations (Levene’s test of homogeneity of variance, p < 0.001).

### Analysis of morphology data

The distribution of different morphological traits (femur length, forewing length and ovipositor length only in females) is shown in **Figure 5**. Since there is a significant difference between the forewing length of males and females (t = −21.331, p < 0.0001), we have treated the data separately by sex. There is no significant difference in the femur length (F = 0.633, p = 0.538 for males; F = 0.134, p = 0.875 for females) in either sex among *L. makaio*, *L. orientalis* and the hybrids. *L. orientalis* crickets have shorter forewings than *L. makaio* (Q = 3.85, p = 0.031 for males; Q = 4.02, p = 0.022 for females) or the hybrids (Q = 4.29, p = 0.015 for males; Q = 5.52, p = 0.002) for either sex. The ovipositor length among the females is also not significantly different among the three groups (F = 2.602, p = 0.093).

**Figure 5:**
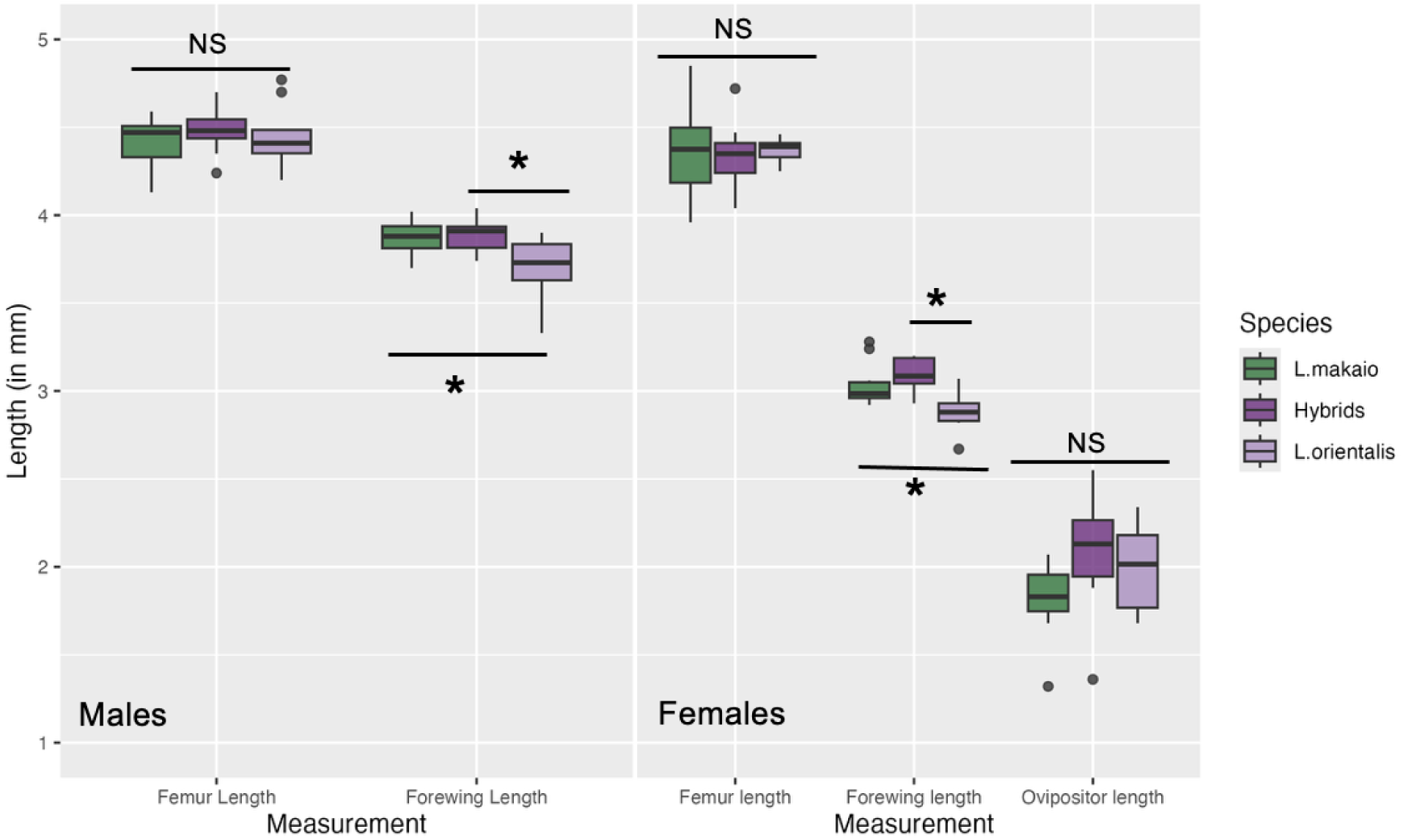
Box plots showing the variation in different morphological traits in males and females of *L. makaio* and hybrids from PW. Hybrid females have significantly longer ovipositors than *L. makaio* females but there is no significant difference in femur or forewing length in either sex between the two groups.

### Population structure analysis

We summarized the genomic variation in our samples in a Principal Components Analysis (PCA) made with genome-wide SNPs. The first two principal components, PC1 and PC2 explain 18.17 % of the total variation (**Figure 6**). Our PCA shows three broad clusters. All the populations from Palikea Peak cluster together but separate by elevation. Starting from site DL (highest elevation site) and SA at the top, where *L. makaio* occurs followed by PW, LS, OK and PB (lowest elevation site) where the hybrids occur. The eight samples of *L. orientalis* from KP are grouped into two clusters containing four samples each. There is some overlap of the SA cluster of *L. makaio* with the hybrid cluster from PW and of the hybrid clusters of OK and PB. Otherwise, all other populations cluster separately. The populations (all Palikea Peak populations except DL) sampled in contemporary times (2021 and 2023) show tighter clustering compared to the historically sampled (2000 and 2001) populations (DL and KP).

**Figure 6:**
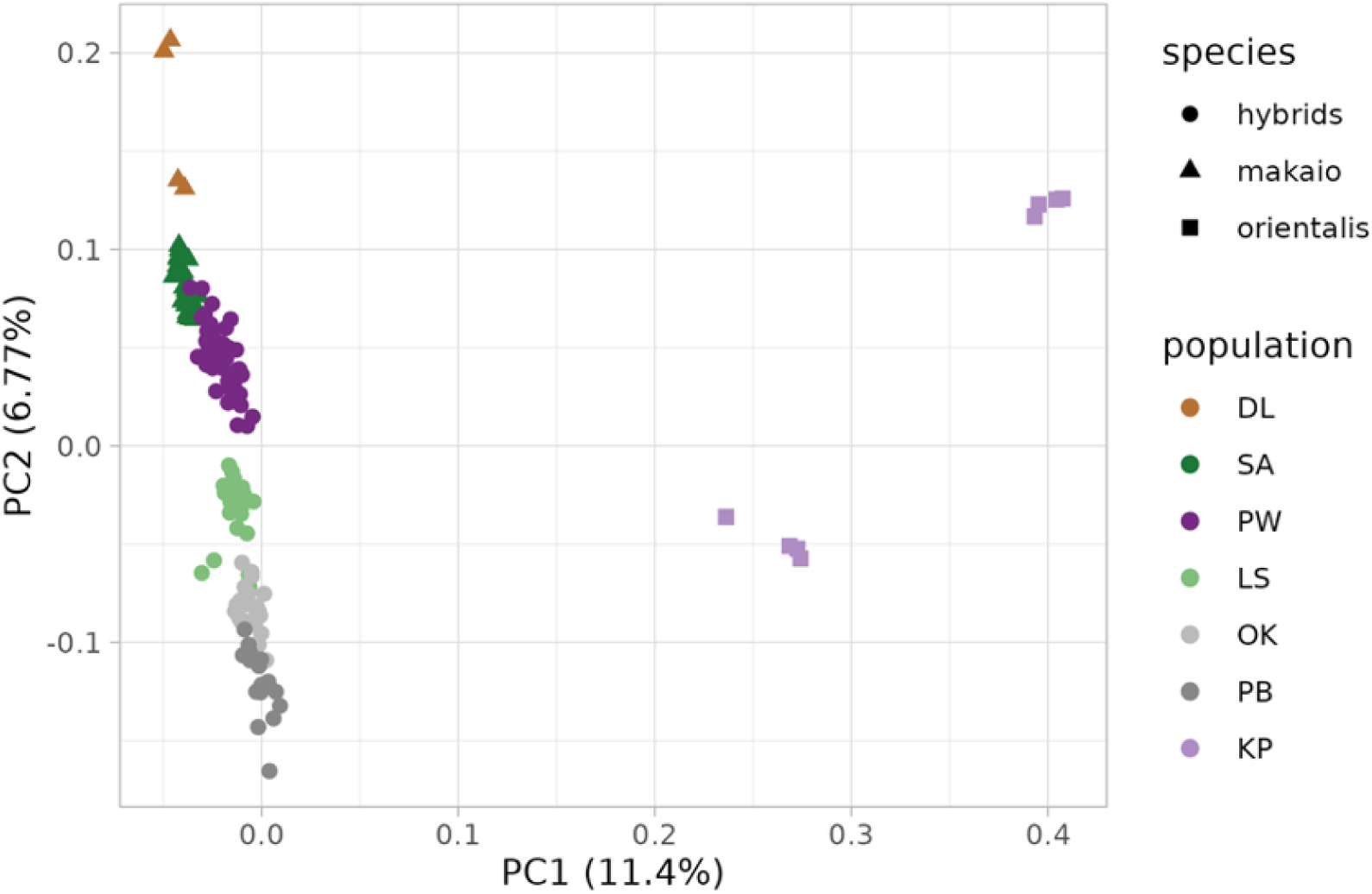
Principal components analyses of 157,419 genome-wide SNPs of 158 samples which included samples from the range of *L. makaio* (DL and SA), *L. orientalis* (KP) and their hybrids (PW, LS, OK and PB). PC axes 1 and 2 explain 11.4% and 6.77% of the variation, respectively.

### Estimation of phylogeny

A maximum likelihood phylogeny of 25 individuals (five each of *L. makaio*, *L. orientalis*, fastest and slowest singers from PW and outgroup *L.eukolea*) with node support values, built in `PhyML 3.0` is shown in **Figure 7**. The schematic in the inset shows a simplified version of the phylogeny. *L.eukolea* is the outgroup and was used to root the tree. Individuals of *L. orientalis* were monophyletic and were grouped closely with the fastest singing hybrids from PW. The fastest and slowest singers from PW and *L. makaio* individuals were paraphyletic. However, the slowest singers from PW grouped closely with *L. makaio*. Slow and fast singers from PW were also grouped closely.

**Figure 7:**
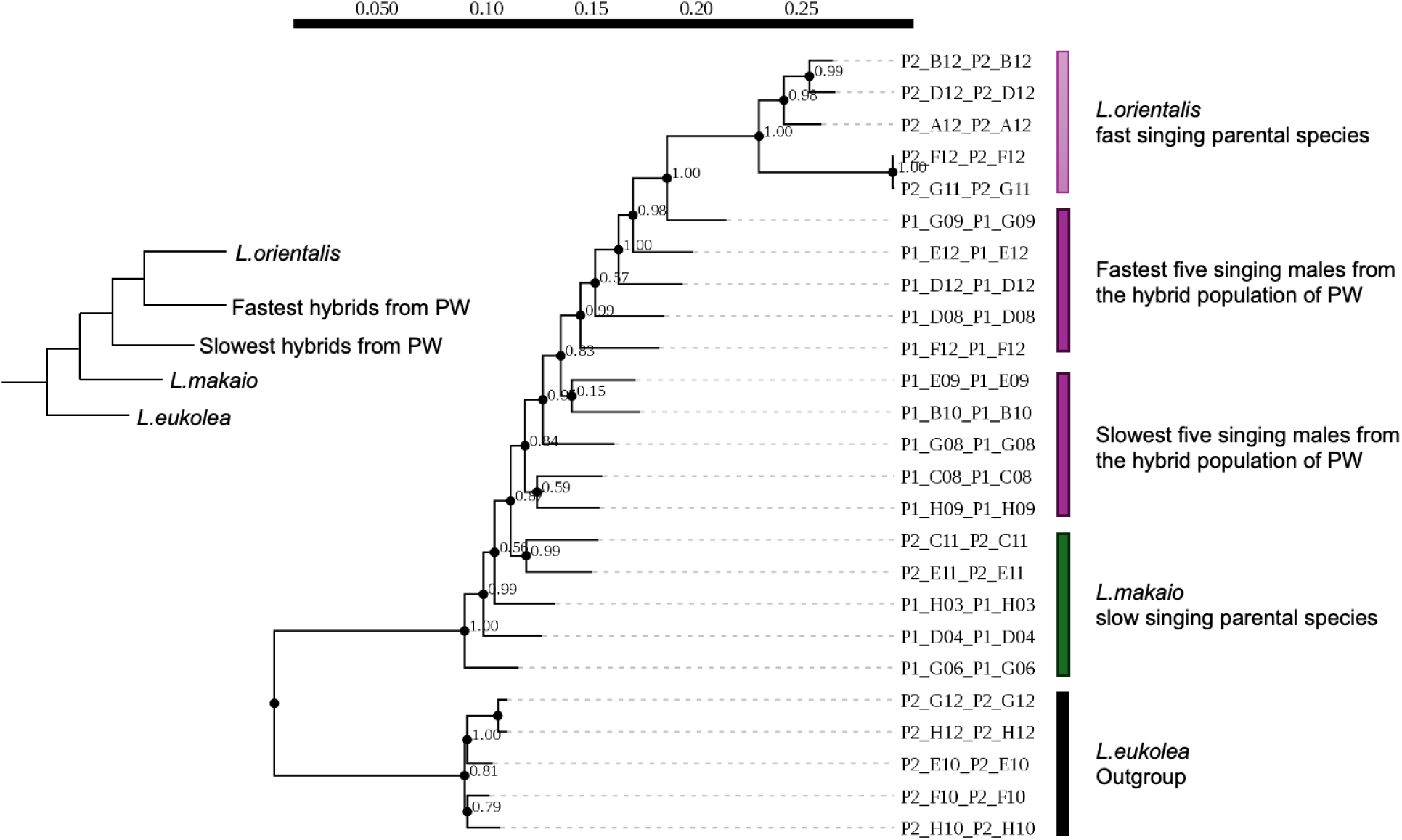
A maximum likelihood phylogeny of 25 individuals built with 48,164 SNPs. The scale bar in the top shows the substitution rate. The numbers near the nodes are branch support values. The tree on the left is a simple schematic version of the actual tree.

### Analysis of hybridization by Triangle Plots

*L. orientalis* (from KP) was chosen as the reference population for calculating hybrid index (HI). So, *L. orientalis* individuals had HI closer to 1, *L. makaio* individuals had HI closer to 0 and hybrids had a range of hybrid indices varying from 0.108 to 0.456 (Mean = 0.264, SD = 0.068). All hybrids had more ancestry of *L. makaio* (HI < 0.5, **Figure 8**). Ancestry of *L. orientalis* decreased as the populations were geographically nearer to where *L. makaio* occurs (SA). For heterozygosity, by the nature of choice of our AIMs, individuals of either parental species are expected to have heterozygosity near zero, and hybrids varied from 0.116 to 0.436 (Mean = 0.22, SD = 0.062; **Figure 9**). A hypothetical F1 hybrid will have a HI of 0.5 (half ancestry from each parent) and a heterozygosity of 1 (all sites are heterozygous because we are using AIMs) and will be placed at the tip of the triangle. A hypothetical BC1 into *L. makaio* (a F1 back-crossed into parental *L. makaio*) will have a HI of 0.25 and a heterozygosity of 0.5. We refer our readers to Figure 1 of Wiens et al., 2025 to have a comprehensive understanding of where different hybrid classes are located on a Triangle Plot. Most hybrids were later generation hybrids with some recent back-crosses into *L. makaio* at PW. There were no F1 hybrids or back crosses to *L. orientalis*. Our results on the classification of hybrids into various classes (following recommendations from Scordato et al., 2017) is summarized in **Table 2**. There is a sharp decrease in heterozygosity from hybrid populations (PW, LS, OK, PB) to parental populations (KP, SA) (One way ANOVA, f-ratio = 24.402, p < 0.00001, **Figure 10**).

**Figure 8:**
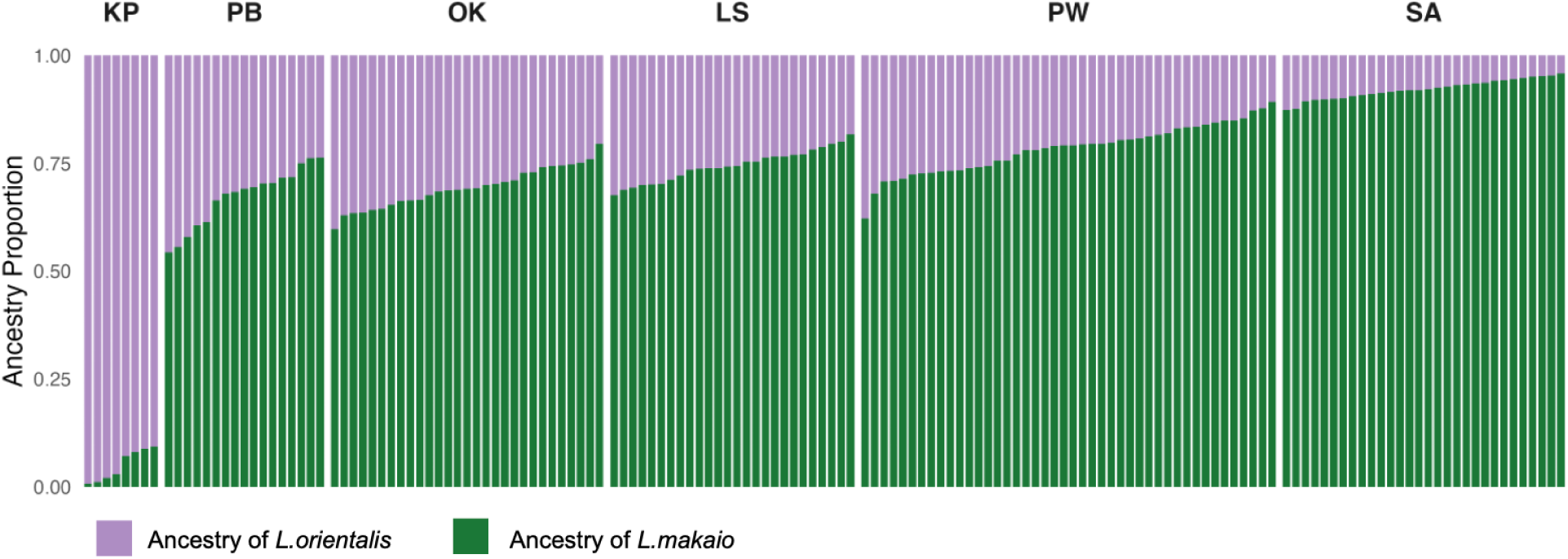
Ancestry proportion (or hybrid index) plot of individuals from each population.

**Figure 9:**
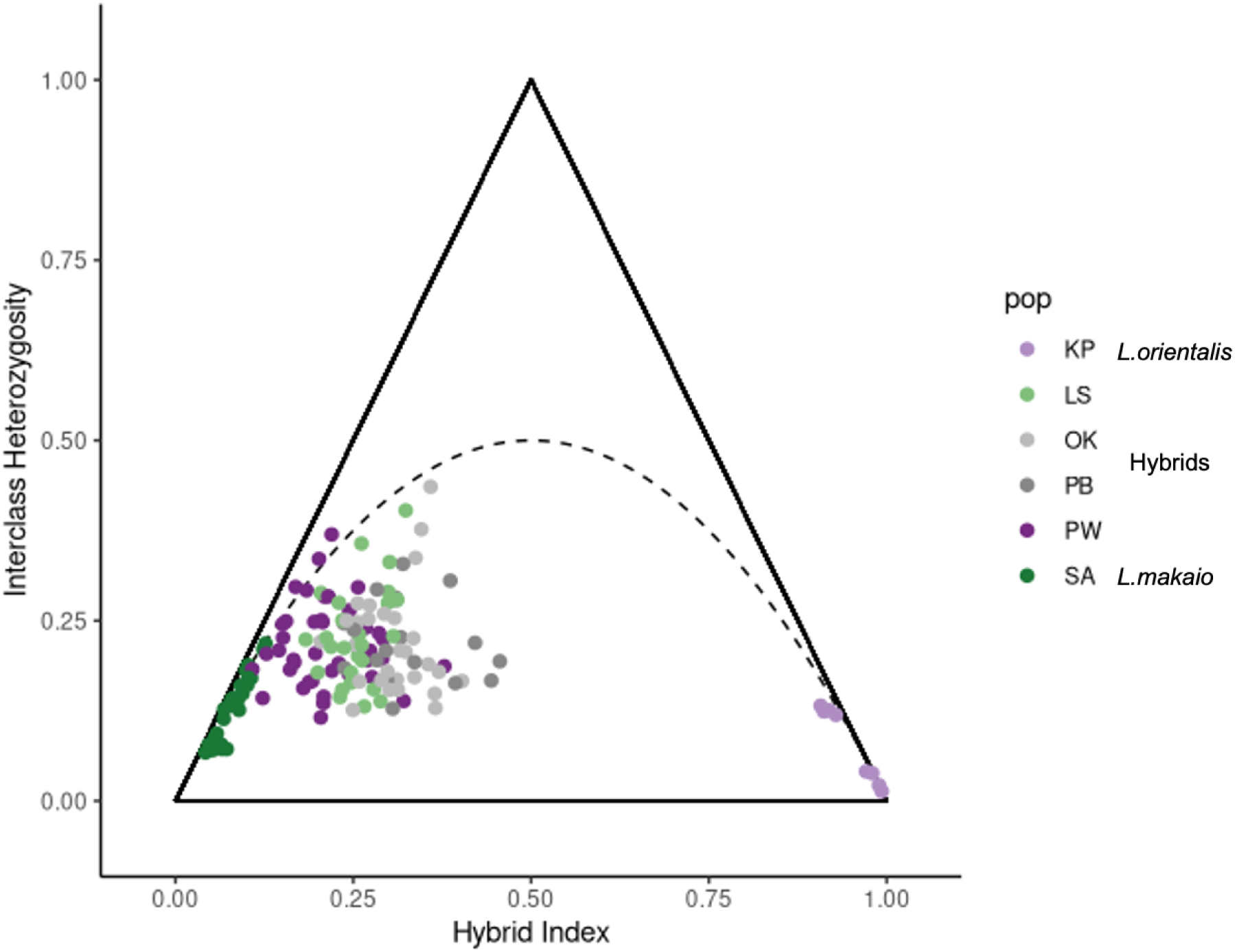
Triangle plot showing genome wide heterozygosity plotted against hybrid index for each individual, colored by population.

**Figure 10:**
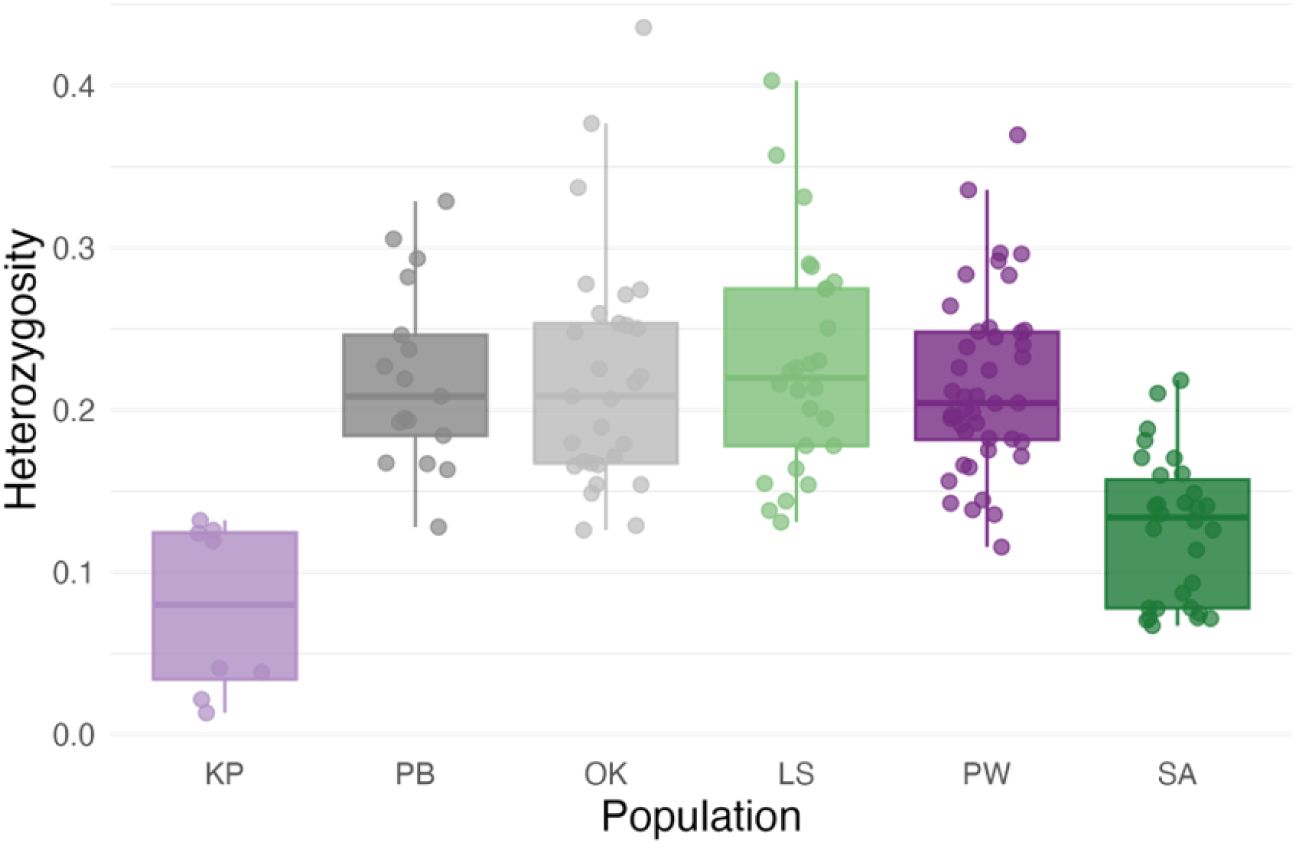
Distribution of heterozygosity scores for each population.

**Table 2:** Distribution of hybrid classes across populations. Crickets with hybrid indices ranging from 0.02 to 0.25 were considered backcrosses to *L. makaio* (BCM). Crickets with hybrid indices >0.25 and <0.75 and heterozygosity ≥ 0.80 were classified as F1 hybrids. Crickets with these intermediate hybrid indices but lower heterozygosity (<0.80) were considered later-generation hybrids (LGH).

| Pop | HI (Mean ± SD) | Heterozygosity (Mean ± SD) | Hybrid class information: Proportion that are later generation hybrids (LGH) or back crosses to *L. makaio* (BCM) |
| --- | --- | --- | --- |
| PW | 0.217 ± 0.057 | 0.215 ± 0.055 | 0.32 LGH, 0.68 BCM |
| LS | 0.256 ± 0.057 | 0.228 ± 0.069 | 0.54 LGH, 0.46 BCM |
| OK | 0.303 ± 0.047 | 0.221 ± 0.073 | 0.9 LGH, 0.1 BCM |
| PB | 0.328 ± 0.069 | 0.22 ± 0.056 | 0.88 LGH, 0.12 BCM |

### Phenotype Genotype Correlations

There is a strong positive and significant correlation (R = +0.74, p = 4.1e-06) between pulse rate and hybrid index (ancestry of the fast singer *L. orientalis*) for the hybrid males (n=29) at PW (**Figure 11**). In other words, hybrid males with a higher ancestry of *L. orientalis*, sing at a faster pulse rate.

**Figure 11:**
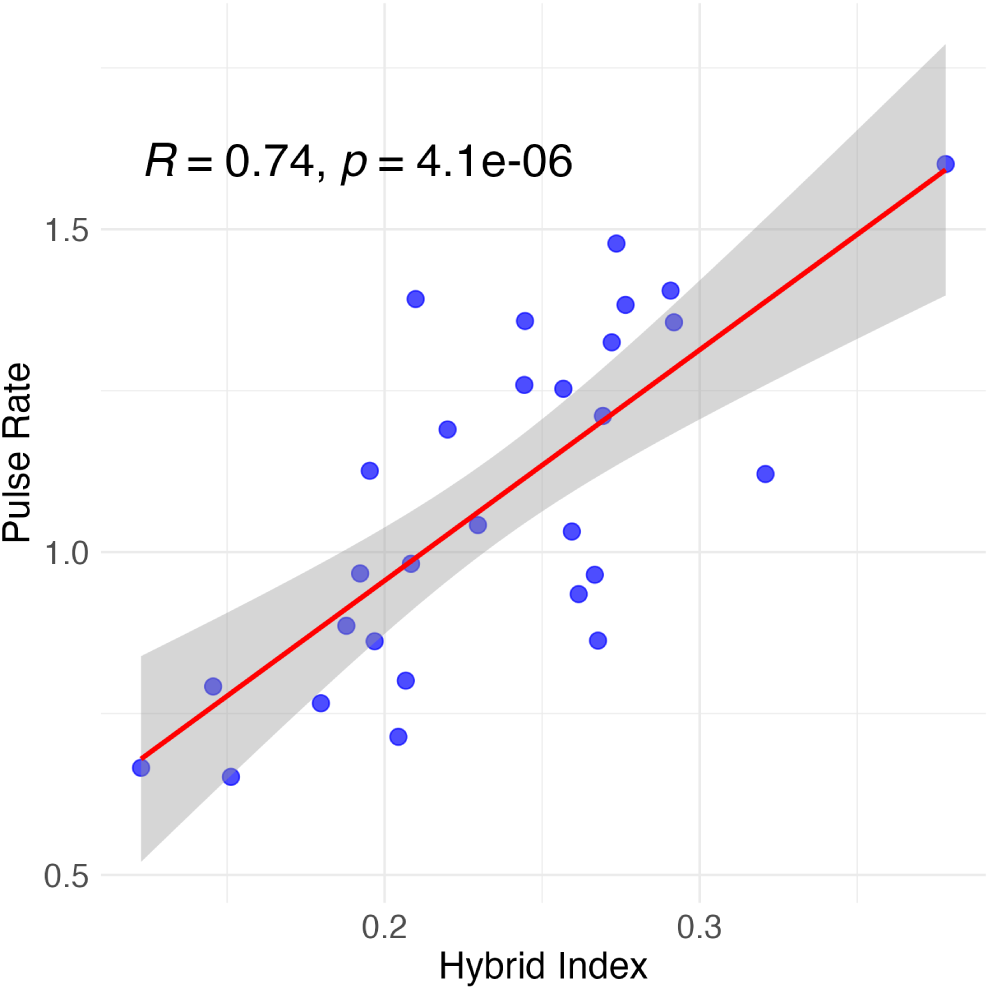
Test of correlation between pulse rate and hybrid index for PW males (n=29).

## Discussion

Recent work has shown that hybridization has emerged as an important process which can fuel species radiations. Documenting natural hybridization in a system is important to support the role of hybridization in accelerating rapid diversification of the lineage. Often, the detection of hybridization in the wild usually begins with an observation of phenotypic intermediacy or admixture of two or more parental phenotypes. This detection is easier for visual phenotypes like wing coloration in insects or plumage patterns in birds but difficult for morphologically cryptic species with non-visual phenotypes that differentiate species. However, the increasing use of next-generation sequencing technologies and genomic analyses have helped the detection of hybrids in cryptic species. In this study, we discovered the first case of natural hybridization in the cryptic Hawaiian crickets of the *Laupala* radiation. Crickets in this genus are morphologically cryptic, but males in different species primarily differ in their songs (pulse rate). The observation of intermediate pulse rates and the huge variation of pulse rates at different sites of a population of crickets inspired our hypothesis of hybridization, which was confirmed with the analyses of genomic data we collected.

### Analysis of pulse rate data

The formal analysis of song data from cricket populations at Palikea Peak confirms our observation in the wild: PW has the highest variance in pulse rate compared to any other sites where hybrids occur (LS, OK, PB). All hybrid pulse rate values are intermediate of that of their parental species, *L. makaio* (at SA) and *L. orientalis* (at KP). Most hybrids have “slower” pulse rates, values which are closer to the slower singing parental species *L. makaio*, compared to the other. This suggests that the hybrids have more of *L. makaio* ancestry, a prediction that was confirmed in our later hybrid index analyses using SNP data. The observation of predominantly slower singing hybrids, combined with the fact that the site where *L. orientalis* occurred in the past (KP) is not found today, suggests that hybridization between parental species is not contemporary. Instead, most hybrids are probably later generation hybrids or back-crosses with *L. makaio*. This prediction was also supported by the Triangle Plot analyses of genomic data.

### Analysis of morphology data

The morphology data confirms our assumption of crypticity. Since we only had some historical samples of *L. orientalis* (from KP) which were used for sequencing, we used wild caught crickets from a different population (MP) for morphometric comparisons. There is no difference in the femur length (of either sex) or ovipositor length (females) among the PW hybrids and the two parental species. However, *L. orientalis* crickets had smaller forewings than either *L. makaio* or the hybrids, but the difference is so tiny (mean difference between male forewing lengths of two parental species is 0.17 mm) that it is still cryptic to the human observer.

### Population structure analysis

We generated a SNP dataset using GBS technology and the first analysis we did was to look at the population genetic structure. The genomic PCA with the first two principal components PC1 and PC2 (which together explains 18.17% of the total variation) clusters the individuals into three broad clusters: one mega cluster of all individuals from Palikea Peak (separated by elevation) and two small clusters of *L. orientalis* from Kipahulu. Instead of clustering together, *L. orientalis* individuals from KP form two clusters, possibly due to strong founder-effects. The *L. orientalis* samples used in this study were historical samples and the two clusters correspond to samples collected in different years (2000 and 2001). Founder-effects are especially strong in island populations because of different waves of colonization and lower effective population sizes (Clegg et al., 2002; Habel & Zachos, 2013; Planes & Lecaillon, 1998). The genomic PCA also roughly corresponds to the geographic distributions of the different cricket populations; as is the case in many genomic PCA studies like the one on European human populations (Novembre et al., 2008), Eurasian barn swallows (Scordato et al., 2017) and a previous work on different populations of *Laupala cerasina* on the big island of Hawaii (Blankers & Shaw, 2024).

### Estimation of phylogeny

If hybridization is happening, we would expect slower hybrids to cluster with the slow parent *L. makaio* and faster hybrids to cluster with the fast parent *L. orientalis*. But the fast and slow hybrids at PW are at the same geographic location and possibly exchanging genes (the genome-wide Fst between the fastest and slowest eight males at PW is 0) so they should cluster together. This is exactly what we found: fastest and slowest singers from PW were paraphyletic, fastest singers grouped closely together with *L. orientalis* and slowest singers grouped closely together with *L. makaio*. We think this is very strong evidence of hybridization as this pattern of phylogenetic relatedness among populations cannot be explained by alternate processes like genetic IBD or local adaptation.

### Analysis of hybridization by Triangle Plots

Hybrid index (HI) and heterozygosity were calculated for all hybrid individuals (and parentals too) at all sites, from ancestry informative SNPs (*L. orientalis* being the reference population for calculation of HI). All hybrids had an excess of *L. makaio* ancestry (maximum HI was 0.436, lower than 0.5) confirming the prediction we had from analyses of pulse rate data alone (abundance of slower hybrids). The ancestry of *L. makaio* decreases (mean HI per population increases) as you move further away from the SA site, an observation which can be predicted from the geographic distribution of the populations alone. A triangle plot of HI vs heterozygosity reveals that there is an absence of recent hybrids, and that most hybrids are later generation hybrids or back crosses with *L. makaio*, with the proportion of backcrosses being higher near the SA site.

### Phenotype Genotype Correlation

A test of correlation between pulse rate and hybrid index for PW showed a strong, positive and significant relationship (R = +0.74, p = 4.1e-06). This is another very important result which confirms that hybridization due to secondary contact is what is causing the huge variation in pulse rates in PW. Even though the spread of hybrid indices is comparable across the different sites at Palikea Peak, other sites (LS, OK and PB) have a very small spread of pulse rates. This impedes us from doing a similar correlation test in other sites due to “restriction of range”: the masking of the stronger relationship between two variables because one variable has very little variance. One might also observe that the mean HI at PW is lower than at other sites (LS, OK and PB) but it still has the highest variance in pulse rate. In other words, crickets at LS, OK and PB have higher mean *L. orientalis* ancestry compared to PW but they don’t sing as fast as PW hybrids. We acknowledge that this is an important observation that warrants further investigation. We speculate that “fast singing” ancestry of *L. orientalis* might be preferentially fixed at/around genes underlying pulse rate modulation in PW hybrids, however this is a very preliminary hypothesis that needs rigorous testing.

Our estimation of phylogeny and results of pulse rate - hybrid index correlation confirms that natural hybridization is indeed occurring in our system. The triangle plot analysis further lets us estimate the generation of hybrids and show that hybridization is older. Although we haven’t performed some classic tests of hybridization like ABBA-BABA test or D-statistics based analyses (mostly because of not having the right populations for those kinds of tests), we think that our analyses were sufficient to demonstrate the occurrence of hybridization. By using a combination of analysis of phenotypic patterns (pulse rate) to generate hypotheses of hybridization and testing those hypotheses with genetic data, we confirm the discovery of the first case of natural hybridization in the genus *Laupala*. Future work will focus on investigating if these hybrid populations are on their path to becoming distinct “hybrid species”. We already know that some of these populations (PB, LS, OK) breed true in the lab (RS,KLS personal observations), which signals that they probably have stabilized hybrid genomes, although this needs to be confirmed with further genomic analyses (Moran et al., 2021; Nevado et al., 2024). We also need to demonstrate reproductive isolation between the hybrid populations and their parental species and demonstrate that this reproductive isolation is caused primarily by hybridization (Schumer et al., 2014).

Our study also paves the way to understand the importance of hybridization in non-adaptive radiations specifically. Oftentimes in adaptive radiations, hybridization helps by aiding a hybrid species to colonize new ecological niches like hybrid *Helianthus* sunflower species colonizing salt-heavy soils (Lexer et al., 2003) or by the spread of useful variants like mimicry rings in *Heliconius* butterflies (Dasmahapatra et al., 2012; Pardo-Diaz et al., 2012; Smith & Kronforst, 2013). In contrast, *Laupala* species are largely ecologically equivalent, raising the question of how hybridization might promote diversification without obvious adaptive benefits. One possibility is that hybridization can create new signal-preference combinations that could drive reproductive isolation through sexual trait divergence. Just like hybrid males sing songs with intermediate pulse rates, likewise hybrid females prefer intermediate pulse rates (Shaw, 2000b) and we have evidence for genomic coupling of male songs and female preferences (Xu & Shaw, 2021). *Laupala* female preference for pulse rate is also unimodal (Shaw, 2000b) and this preference can drive the stabilization of pulse rate values in a new hybrid population. A non-adaptive radiation like *Laupala* would be the perfect “model clade” to pursue these questions and to improve our understanding of the role of hybridization in fueling explosive species radiations.

## Acknowledgements

We are grateful to Ron and Linda Nagada for hosting us while conducting fieldwork on the island of Maui, Hawaii. We thank Bhaavya Srivastava for helping with fieldwork and animal care. We thank the entire Shaw lab and the “Lunch Bunch” discussion group in the Department of Neurobiology and Behavior at Cornell University for fruitful discussions on the project. We thank Niko Hensley for giving feedback on the manuscript.

## Data and Code availability

All data and R codes will be made publicly available upon publication of the manuscript.

## Funding

RS was supported by the Sustainable Biodiversity fund from Cornell Atkinson Center for Sustainability, Research Travel Grant from Cornell Graduate School, Theodore J. Cohn research fund from the Orthopterists’ Society, Student Research Grant from the Animal Behavior Society, R.C. Lewontin Early Award from the Society for the Study of Evolution, research grants from the Cornell chapter of Sigma Xi and departmental research grants from the Department of Neurobiology and Behavior. KLS was supported by funds from the NSF grant IOS-2128521.

## Notes

### Competing Interest Statement

The authors have declared no competing interest.

