## Supplementary Information for "Genomic data confirms phenotypic predictions of hybridization between cryptic Hawaiian cricket species"

**Table S1:** GPS coordinates and elevation of each collection site with the number of crickets sequenced from each site. We collected from sites SA, PW, LS, OK, PB and MP on 2021 and 2023 and used frozen samples from past collections at DL and KP.

| Collection site | GPS coordinates | Elevation (m) | Number of samples sequenced |
|---|---|---|---|
| SA | 20.6764, -156.0728 | 681.12 | 30 |
| PW | 20.6762, -156.0712 | 676.16 | 44 |
| LS | 20.6743, -156.0699 | 604.36 | 26 |
| OK | 20.6718, -156.0671 | 456.43 | 29 |
| PB | 20.6699, -156.065 | 409.04 | 17 |
| MP | 20.804, -156.1085 | 452.35 | 19 |
| DL | 20.704, -156.1167 | - | 4 |
| KP | 20.670, -156.054 | - | 8 |

### Isolation by distance at Palikea Peak

**Table S2:** Distance (unshaded) and Fst values (shaded) between populations at Palikea Peak

| | SA | PW | LS | OK | PB |
|---|---|---|---|---|---|
| SA | | 0.002142 | 0.0056749 | 0.011123 | 0.015485 |
| PW | 168.003207 | | 0.00091495 | 0.0053443 | 0.007239 |
| LS | 389.145227 | 260.922928 | | 0 | 0 |
| OK | 814.696001 | 685.281609 | 428.973125 | | 0 |
| PB | 1120.24295 | 989.054456 | 733.099051 | 307.573308 | |

**Figure S1:** A plot of pairwise Fst vs geographic distance shows a clear pattern of isolation by distance

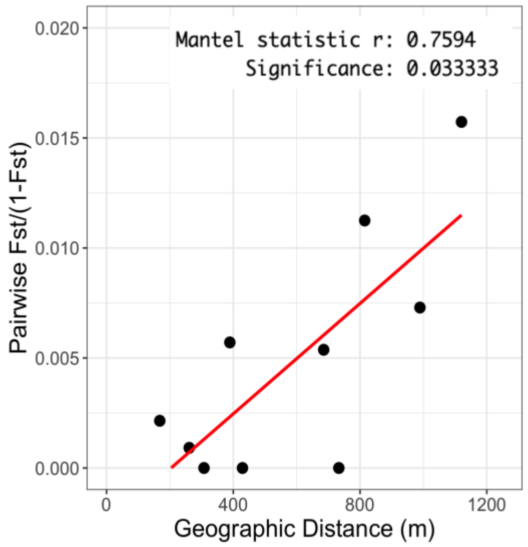
